# Intestinal dysbiosis in carbapenem-resistant Enterobacteriaceae carriers

**DOI:** 10.1101/855718

**Authors:** Hila Korach-Rechtman, Maysaa Hreish, Carmit Fried, Shiran Gerassy-Vainberg, Zaher S Azzam, Yechezkel Kashi, Gidon Berger

**Affiliations:** Faculty of Biotechnology and Food Engineering, Technion–Israel Institute of Technology, Haifa, Israel; Ruth & Bruce Rappaport Faculty of Medicine, Technion - Israel Institute of Technology, Haifa, Israel; Department of Internal Medicine ‘B’, Rambam Health Care Campus, Haifa, Israel

**Keywords:** Carbapenem-resistant Enterobacteriaceae (CRE), Microbiome, Intestinal dysbiosis, antibiotic resistance

## Abstract

Infection with Carbapenem-Resistant Enterobacteriaceae (CRE) became an important challenge in health-care settings and a growing concern worldwide. Since infection is preceded by colonization, an understanding of the latter may reduce CRE-infections. We aimed to characterize the gut microbiota after colonization by CRE, assuming that an imbalanced gastrointestinal tract (GIT)-associated microbiota precedes CRE-colonization.

We evaluated the GIT-microbiota using 16S rRNA genes sequencing extracted of fecal samples, collected from hospitalized CRE-carriers, and two control groups of hospitalized non-carriers and healthy adults. The microbiota diversity and composition in CRE-colonized patients differed from that of the control groups participants. These CRE-carriers displayed lower phylogenetic diversity and dysbiotic microbiota, enriched with members of the Enterobacteriaceae family. Concurrent with the bloom in Enterobacteriaceae, a depletion of anaerobic commensals was observed. Additionally, changes in several predicted metabolic pathways were observed for the CRE-carriers. Concomitant, we found higher prevalence of bacteremia in the CRE-carriers.

Several clinical factors that might induce change in the microbiota were examined and found as insignificant between the groups.

CRE-colonized patients have dysbiotic gut microbiota in terms of diversity and community membership, associated with increased risk for systemic infection. Our study results provides justification for attempts to restore the dysbiotic microbiota with probiotics or fecal transplantation.

## Background

The emergence and spread of highly antibiotic-resistant bacteria represent a major clinical challenge. In recent years, the numbers of infections caused by bacteria such as *Clostridium difficile*, Methicillin-resistant *Staphylococcus aureus*, and Vancomycin-resistant Enterococcus, have increased markedly [1]. Carbapenem-resistant Enterobacteriaceae (CRE) are highly drug-resistant pathogens with a rapidly increasing incidence in a variety of clinical settings [2].

Infections caused by CRE have been associated with increased cost and length of stay as well as frequent treatment failures, and death [2]. There are several known risk factors [2,3], including CRE-carriage in the GIT, since it serves as a source for subsequent clinical infection in approximately 9% of carriers [4]. Moreover, CRE-carriers serve as a major reservoir for its dissemination in healthcare facilities [4,5].

The complex commensal microbiota that normally colonizes mucosal surfaces in healthy individuals allows colonization resistance and inhibits expansion and domination by antibiotic-resistant exogenous bacteria such as Enterobacteriaceae [6,7]. Microbial dysbiosis may lead to an overgrowth of antibiotic resistant pathogens [8], which can be calamitous for susceptible patients, resulting in bacteremia and sepsis [9] and is associated with increased risk for transmission due to increased shedding to the environment [10,11].

It is reasonable to assume that alteration of the normal microbiota may be associated with the development of CRE-carriage. Therefore, we aimed to determine the structure of GIT-microbiota in CRE-colonized patients.

## Materials and Methods

### Study design and participants

The study population was comprised of two control groups (15 healthy adults and 22 hospitalized non-CRE-carriers), and one group of CRE-carriers (*n*=40). Hospitalized adults were recruited from the Division of Internal Medicine at Rambam Health Care Campus (Israel); non-hospitalized healthy participants were recruited from the local community, and most likely were not CRE-carriers. Within the hospitalized groups, distinction of CRE-carriage was established upon hospitalization and every other week by routine screening for rectal carriage.

The following exclusion criteria were applied to the healthy group to avoid factors capable of altering the microbiome: current smoker, active or recent (6 months) chemotherapy and/or radiation treatment, homeopathic preparations user, currently suffering any infectious disease, chronic or acute gastro intestinal track disease including *Clostridium difficile* infection, and/or antibiotic treatment or vaccination within the last 6 months prior to sample collection. This group individuals volunteered and were recruited randomly by hospital staff. Nonetheless, due to the lack of antibiotic treatment, difference in participant’s average age and the fact they were not hospitalized under similar conditions, the majority of comparative analyses were conducted between the two hospitalized groups.

Both hospitalized groups included in this study were hospitalized for at least 7 days at the same healthcare facility. Inclusion criteria included patient hospitalized for at least 7 days, which receive minimum 7 days of antibiotic treatment.

Written informed consent was obtained from all study participants. The study was approved by the Rambam Health Care Campus ethics committee (approval number 0418-14-RMB). This study conformed to the Helsinki Declaration and to local legislation.

Clinical analyses were performed on the entire study population (*n*=77), and 55 samples from the three groups were sent for sequencing (the earliest recruits). Clinical variables including medical background, and laboratory parameters were retrieved from medical records (see table 1 and 2).

**Table 1:**
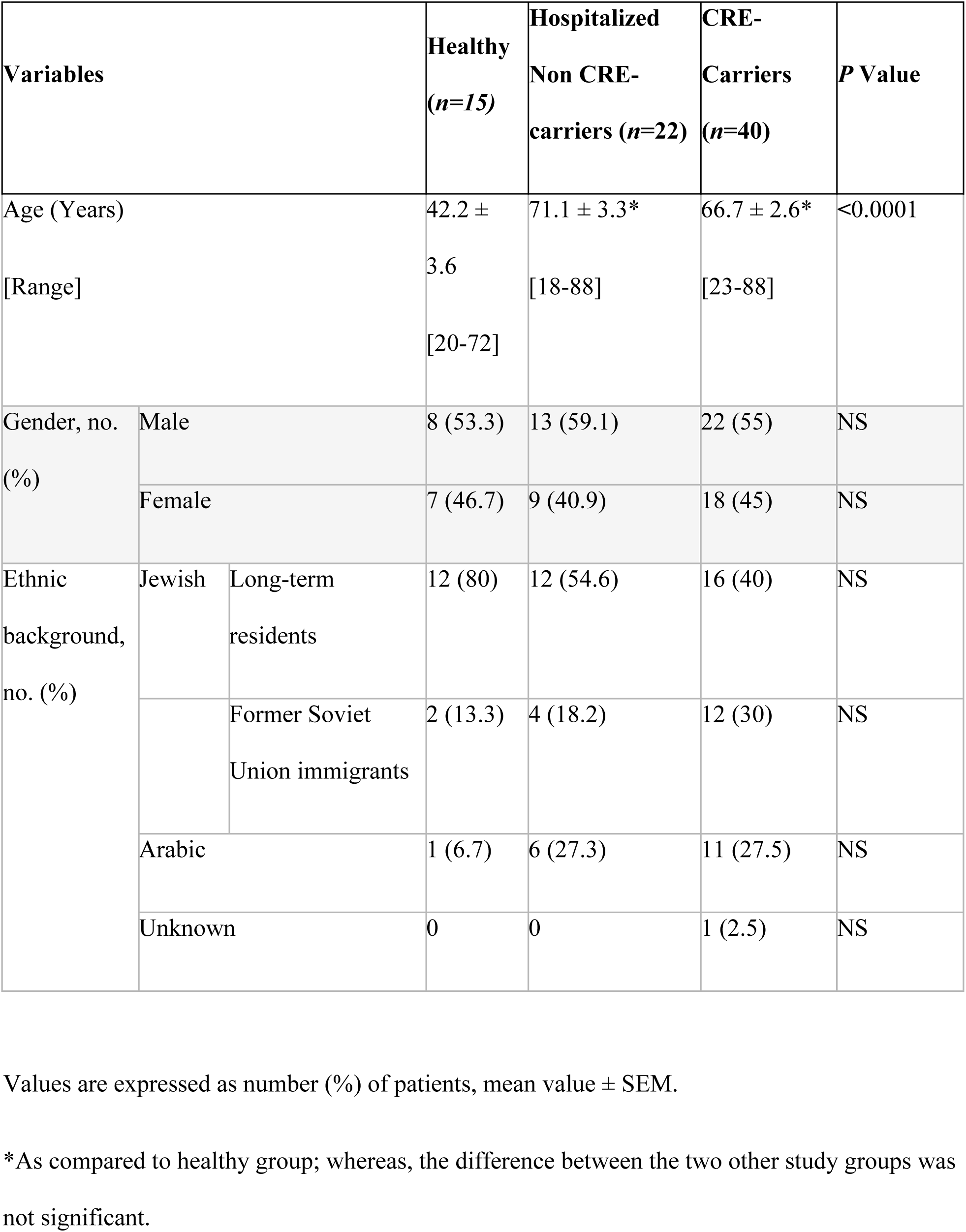
Demographic characteristics of the study cohort.

**Table 2:**
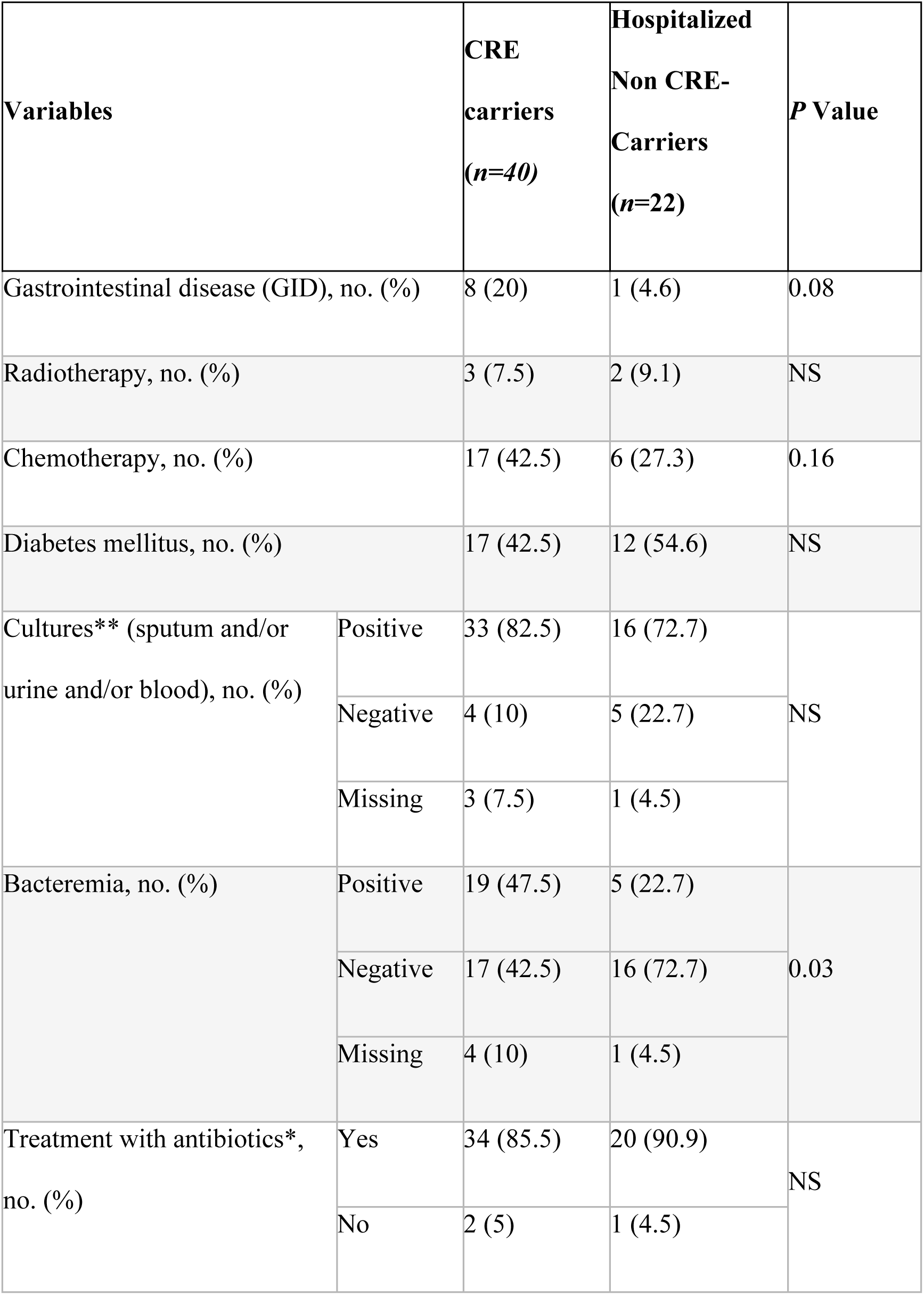
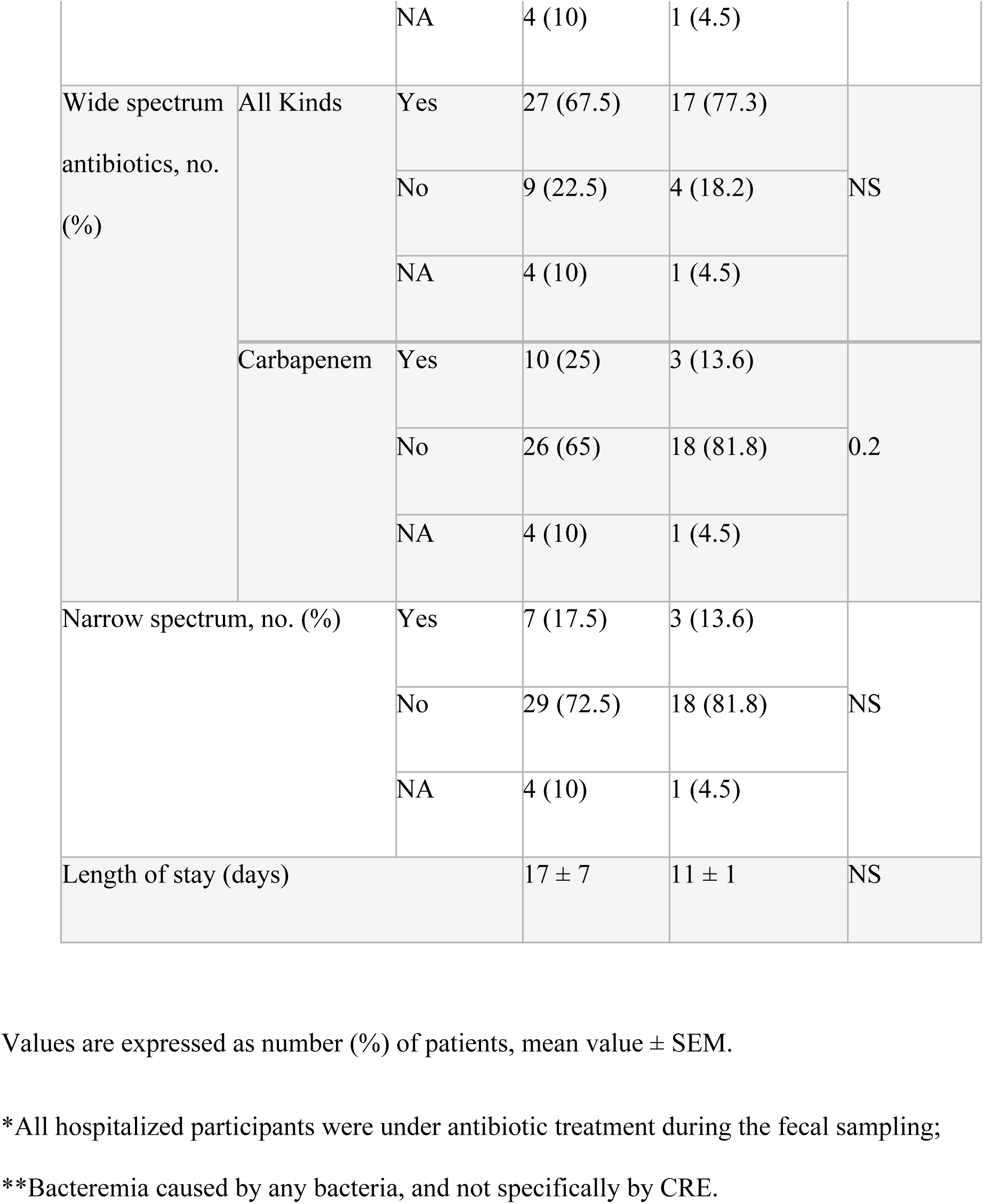
Clinical characteristics of the hospitalized study cohort.

| **Variables** |  | **CRE carriers**<br>**(n=40)** | **Hospitalized**<br>**Non CRE-Carriers**<br>**(n=22)** | **P Value** |
| --- | --- | --- | --- | --- |
| Gastrointestinal disease (GID), no. (%) |  | 8 (20) | 1 (4.6) | 0.08 |
| Radiotherapy, no. (%) |  | 3 (7.5) | 2 (9.1) | NS |
| Chemotherapy, no. (%) |  | 17 (42.5) | 6 (27.3) | 0.16 |
| Diabetes mellitus, no. (%) |  | 17 (42.5) | 12 (54.6) | NS |
| Cultures** (sputum and/or urine and/or blood), no. (%) | Positive | 33 (82.5) | 16 (72.7) | NS |
|  | Negative | 4 (10) | 5 (22.7) |  |
|  | Missing | 3 (7.5) | 1 (4.5) |  |
| Bacteremia, no. (%) | Positive | 19 (47.5) | 5 (22.7) | 0.03 |
|  | Negative | 17 (42.5) | 16 (72.7) |  |
|  | Missing | 4 (10) | 1 (4.5) |  |
| Treatment with antibiotics*, no. (%) | Yes | 34 (85.5) | 20 (90.9) | NS |
|  | No | 2 (5) | 1 (4.5) |  |

|  |  |  |  |  |  |
| --- | --- | --- | --- | --- | --- |
|  |  | NA | 4 (10) | 1 (4.5) |  |
| Wide spectrum antibiotics, no. (%) | All Kinds | Yes | 27 (67.5) | 17 (77.3) | NS |
|  |  | No | 9 (22.5) | 4 (18.2) |  |
|  |  | NA | 4 (10) | 1 (4.5) |  |
|  | Carbapenem | Yes | 10 (25) | 3 (13.6) | 0.2 |
|  |  | No | 26 (65) | 18 (81.8) |  |
|  |  | NA | 4 (10) | 1 (4.5) |  |
| Narrow spectrum, no. (%) |  | Yes | 7 (17.5) | 3 (13.6) | NS |
|  |  | No | 29 (72.5) | 18 (81.8) |  |
|  |  | NA | 4 (10) | 1 (4.5) |  |
| Length of stay (days) |  |  | 17 ± 7 | 11 ± 1 | NS |

Values are expressed as number (%) of patients, mean value ± SEM.
\*All hospitalized participants were under antibiotic treatment during the fecal sampling;
\*\*Bacteremia caused by any bacteria, and not specifically by CRE.

### Sample collection

Fresh fecal samples were collected from hospitalized participants by the research cadre (CRE-carriers and non-carrier groups). Following collection, swabs were stored at −80^º^C. Samples from healthy participants were self-collected by participants, and transported in a freezer pack to the laboratory within 24 hours of collection, and then stored at −80^º^C.

### Microbiota sequencing and taxonomy assignment

Total DNA was extracted from the samples using QIAamp Fast DNA Stool Mini Kit (QIAGEN Inc., Hilden, Germany), according to manufacturer`s instructions. Genomic DNA was PCR (Polymerase Chain Reaction) amplified using primers CS1-515F (5’-GTGCCAGCMGCCGCGGTAA) and CS2-806R (5’-TACGGTAGCAGAGACTTGGTCTGGACTACHVGGGTWTCTAAT) targeting the V4 region of the 16S rRNA gene. Samples were sequenced at the DNA Services (DNAS) Facility, Research Resources Center (RRC), University of Illinois at Chicago (UIC), using dual PCR strategy [1,2]. PCR amplifications were performed in 25 µl reactions. Each PCR mixture contained 0.2 mM deoxynucleoside triphosphates, 0.4 μM forward and reverse primers, 0.02 U of Taq polymerase (SuperNova, JMR Holding, Kent) per μl, 1x reaction buffer (containing 1.5 mM MgCl2) and 10-100 ng of genomic DNA. The reactions were carried out in a Veriti® 96-Well Thermal Cycler (Applied Biosystems) as follows: 95°C for 5 minutes, followed by 28 cycles of 95°C for 30 sec, 50°C for 30 sec and 68°C for 30 sec and a final elongation step of 5 min at 68°C. PCR amplification products were verified by gel (1.2% Agar) electrophoresis and visualized by UV fluorescence. A second PCR amplification (eight cycles) was performed at the DNAS facility (UIC) to incorporate barcodes and sequencing adapters into the final PCR products. The primers for the second PCR reactions included Illumina sequencing adapters, sample-specific barcodes and CS1 and CS2 linkers (Fluidigm) [3]. Sequencing was performed using Illumina MiSeq platform at the DNAS facility using standard V3 chemistry with paired-end, 300bp reads. Sequences were automatically binned according to multiplex barcodes on instrument, as described previously [3].

Sequenced data were processed using QIIME 1.8.0 (Quantitative Insight into Microbial Ecology) pipeline [12]. Operational taxonomic units (OTUs) were defined based on 97% similarity clustering using uclust. Taxonomy was assigned against Greengenes database (v13_8) as reference [13]. Diversity analyses were calculated with a rarefied OTU table containing 40,000 reads per sample, and were conducted twice, with and without taxa belonging to the Enterobacteriaceae family.

### Microbiota composition and metabolic analysis

Diversity analyses were calculated with a rarefied OTU table containing 40,000 reads per sample.α-diversity was calculated using the Shannon diversity index and visualized by box-plots using the statistical software environment R

β-diversity was determined by computing weighted UniFrac distance, and the resulting matrices were visualized by Principal Coordinates Analysis (PCoA) plots. The PCo score comparison based on the Kruskal-Wallis test was used to test for differences in community composition among the study groups and the clinical variables using R.

Linear discriminant analysis coupled with effect size measures (LefSe) [29] was used to identify the taxa differentiating the experimental groups. Taxa showing LDA<2.5 were further compared to the clinical variables and to the different antibiotic treatments using R.

Bacterial metabolic activity abundance, as defined by the Kyoto Encyclopedia of Genes and Genomes (KEGG) [30], were generated by Phylogenetic Investigation of Communities by Reconstruction of Unobserved States (PICRUSt version 1.1.3) [31], using closed reference OTUs picked by QIIME over the same set of sequences. LefSe was used to identify functional attributes differentiating between CRE-carriers and non-carriers groups. Selected functional attributes were compared based on the Kruskal-Wallis test using R. Relative abundance of bacteria, with an LDA>2 in the LefSe analyses between hospitalized CRE-carriers and non-carriers was correlated with level 2 and 3 (L2, L3) functional profiles. Pearson correlation coefficients and FDR corrected p-values were calculated using R.

## Results

To study the microbiota profile in CRE-carriage we analyzed the clinical parameters and microbial composition of three groups: hospitalized CRE-carriers, hospitalized non-carriers, and healthy controls.

### Study cohort clinical characteristics

The demographic and clinical characteristics of the study cohort for all groups are presented in Tables 1 and 2. There were no significant differences between the groups regarding confounding factors such as gender, ethnic origin, gastrointestinal disease, radiotherapy, chemotherapy, and diabetes mellitus. Moreover, the comparison between these factors and the microbial profile described below was insignificant.

Hospitalized patients were older than the healthy individuals (average ages 68.3 and 42.2, respectively). In general, antibiotic usage (broad vs. narrow spectrum), and positive culture prevalence were similar in both hospitalized groups. However, the rate of bacteremia was twice as high in the CRE-carriers.

Regarding antibiotics usage, it was noted that Vancomycin and Tazocin treatments were used more in the CRE-carriers, while Amikacin and Rocephin treatments were used more in the hospitalized non-carriers (Kruskal-Wallis *p*<0.05). Notably, most comparative analyses conducted between the two hospitalized groups (excluding the healthy group), because of the different age average, the strict exclusion criteria and the lack of antibiotic treatment, which effect the microbiota.

The prevalence of positive urine, sputum, and blood cultures (not specifically by CRE) was around 80% and similar between both hospitalized groups. However, a higher bacteremia rate was found in the in the CRE-carriers as compared to the non-carriers (53% vs 24%, respectively, *p*=0.03). Interestingly, 74% (14 out of 19) of the bacteremias detected in the CRE-carriers were caused by Enterobacteriaceae, of which 57% had *Klebsiella pneumoniae*; only one patient had bacteremia with *K. pneumoniae*, a CRE due to the *K. pneumoniae* carbapenemase [KPC]); five patients had extended spectrum beta-lactamase (ESBL) *K. pneumoniae*, and two had *K. pneumoniae*.

### Microbiota characterization

To characterize the microbiota, participants fecal DNA was subjected to 16S rRNA genes sequencing.

Taxonomic classification revealed that the dominant bacterial phyla were Bacteroidetes (56-62%), Firmicutes (19-35%), and Proteobacteria (6-21%) (Fig 1A). Firmicutes prevalence was significantly lower in the CRE-carriers compared to non-carriers and healthy controls (*p*<0.005); Proteobacteria were significantly higher in the CRE-carriers (*p*<0.005). The ratio between Firmicutes to Bacteroidetes, considered highly relevant in human gut microbiota composition, was the lowest in the CRE-carriers (0.35±0.05), higher for hospitalized non-carriers (0.41±0.05), and highest in the healthy group (0.63±0.05).

**Fig 1.**
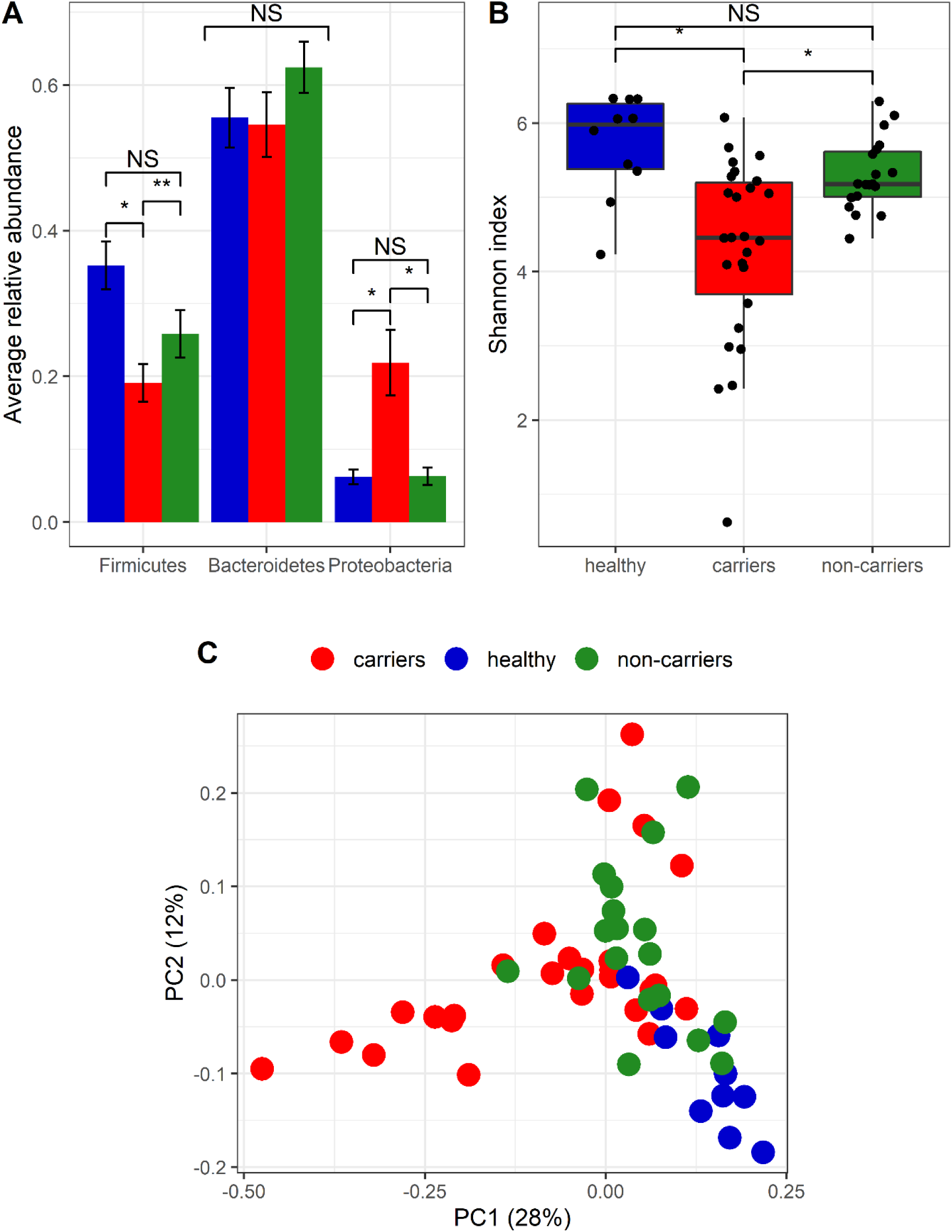
Microbiota composition in the healthy, hospitalized CRE carriers and non-carriers. Bacterial composition was assessed by Illumina MiSeq 16S rRNA genes sequencing of fecal DNA samples. (A) The three dominant phyla relative abundances in the three experimental groups. (B) α-diversity between microbial communities were box-plotted based on Shannon diversity index. (C) β-diversity between microbial communities were clustered using Principal Coordinates Analysis (PCoA) based on weighted UniFrac measure. \**p*<0.005, \*\**p*<0.001; Healthy group: blue dots; CRE carriers: red dots; non-carriers: green dots.

### Microbial diversity and composition

Microbial richness assessment determined using the Shannon index, revealed that CRE-carriers had significantly lower richness as compared to the other groups (*p*<0.005) (Fig 1B). Interestingly, the healthy and non-carriers groups did not differ in the richness measure.

The bacterial communities of the three groups were compared using weighted UniFrac based on PCoA (Fig 1C). The samples from healthy individuals clustered separately from hospitalized participants. The PC1 and PC2 vectors significantly discriminated between the groups (Kruskal-Wallis *p*<0.001).

LefSe was used to identify bacterial taxa associated with CRE-carriage, by comparing the microbiota of CRE-carriers and non-carriers (Fig 2A). The CRE-carriers had a significantly increased prevalence of different genera belonging to the Enterobacteriaceae family including *Pantoea, Enterobacter*, *Klebsiella* and *Erwinia*. Decreased prevalence was observed for the Rikenellaceae, Barnesiellaceae, and genera belonging to Clostridiales including *Ruminococcus*, *Faecalibacterium*, *Coprococcus*, and *Ornithobacterium* (see LDA scores in Fig 2C), an anaerobic commensals, with potentially important beneficial roles for the host [6,7,12]. The decreased abundance of the Barnesiellaceae family among CRE-carriers, was also significant in analysis of the microbiota with the removal of all OTUs belonging to Enterobacteriaceae family.

**Fig 2:**
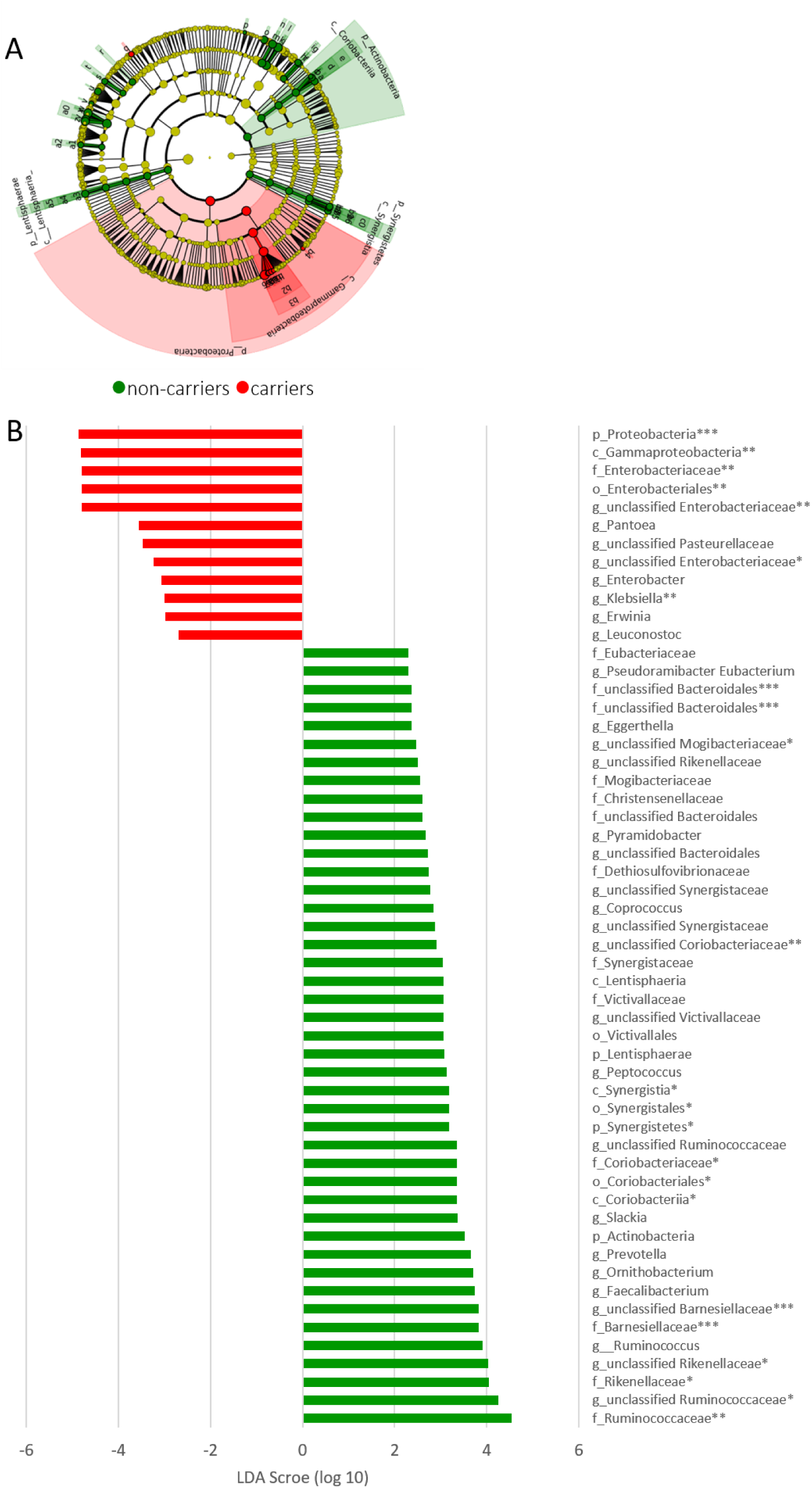
Bacterial markers associated with CRE carriers. Differentially abundant Operational Taxonomic Units (OTUs) between CRE carriers (red) and non-carriers (green) were identified using LefSe and presented by (A) Cladogram; (B) Histogram of LDA scores (log 10). Groups defined according to the microbiota origin. Only taxa meeting LDA ≥ 2 and *p*<0.05 are shown. \**p*<0.01, \*\*\**p*<0.005, \*\*\**p*<0.001.

### Functional prediction of the microbiota

In this study, we used 16S rRNA genes to study microbial communities in CRE-carriage. However, this marker gene cannot directly identify metabolic or other functional capabilities of the microorganisms. Nonetheless, PICRUSt is a technique that uses evolutionary modeling to predict metagenomes from 16S data and a reference genome database. We used this nd their correlation with relative abundances of selected bacteria identified by LefSe.

We found that CRE-carriers were enriched in functional categories associated with xenobiotics biodegradation and metabolism (L2), and amino benzoate degradation (L3) (LDA=3.03, 2.14 respectively; *p*<0.02; Fig S2). Moreover, Enterobacteriaceae family positively correlated with xenobiotics biodegradation and metabolism (R=0.534; FDR *p*<0.003). Other functional categories changed in CRE-carriers included reduction in histidine metabolism (L3), elevation in ubiquinone and other terpenoid-quinone biosyntheses (L3), and tryptophan metabolism (L3) (Fig S1).

## Discussion

This study has demonstrated that CRE-colonized patients have dysbiotic microbiota in terms of community membership with different functional metabolic microbiota profiles.

The intestinal microbiota can protect itself against colonization with new bacteria (colonization resistance), while dysbiosis is exploited, apparently, by CRE for colonization. On the other hand, it is also possible that the established CRE-colonization induce significant perturbations to the microbiota, which in turn may act as a pathogenic community to perpetuate host pathology [13].

We observed that healthy individuals has higher microbial diversity while CRE-carriers has the lowest diversity (Fig 1B) [14,15]. Moreover, we observed three clusters indicating different microbial community structure for each of the experimental groups (Fig 1C). This is in agreement with numerous studies that have shown reduced bacterial diversity in obesity, inflammatory bowel disease, irritable bowel syndrome, and type-2 diabetes mellitus [14,15].

Few specific taxa of the microbiota were different between the CRE-carriers and non-carriers. First, in addition to the CRE-itself, we observed increased Enterobacteriaceae abundance (*Enterobacter*, *Erwinia*, *Pantoea*, *Klebsiella*), among which were resident species with virulence potential that are normally kept at low levels. This consequently predisposes the host to infections with life-threatening sequelae caused by the CRE-itself and potentially other pathobionts.

Second, concurrently with the bloom in Enterobacteriaceae, a depletion of anaerobic-commensals was observed (Fig 2), among which were *Coprococcus* and *Faecalibacterium* abundance, an important short-chain fatty acids (SCFA) producing commensal bacteria [12]. These SCFAs are physiological byproducts of carbohydrate fermentation by the microbiota, and serve to salvage energy for the host, enhance the mucosal barrier, inhibit intestinal inflammation, and oxidative stress [16]. Among the functional consequences of reduction in anaerobic-bacteria, is a reduced metabolic capacity, often exemplified by a decline in SCFAs production. Dysbiosis caused by broad-spectrum antibiotics (e.g. clindamycin, cephalosporins), which in our case can presumably enable CRE-colonization, is commonly associated with low intestinal SCFAs levels [16].

Colonization resistance depends on microbiota diversity, as well as microbial composition. The intestinal microbiota can protect efficiently against colonization by many enteric pathogens. Therefore, it is not surprising that during dysbiosis, intestinal colonization resistance is impaired. Interestingly, it was shown that *Barnesiella* spp., which in our study was less abundant in CRE-carriers as compared to non-carriers, has the ability to restrict the growth of intestinal pathogens, limit colonization with highly antibiotic-resistant bacteria, and is required to prevent expansion of oxygen-tolerant bacteria such as Enterobacteriaceae [6,7].

As an outcome of dysbiosis, predictions of metabolic function also indicated a profile shift. In CRE-carriers we found changes in abundance of several pathways including increased histidine metabolism, and decreased ubiquinone and other terpenoid-quinone biosynthesis, tryptophan metabolism, xenobiotics biodegradation and metabolism, and amino benzoate degradation (Fig S1). These metabolic alterations have been previously linked to modulation of the immune system response to pathogens and the adaptive immune system activation [17,18], and compromised intestinal epithelial barrier and function, which allow bacterial translocation [6,19].

Xenobiotics biodegradation and metabolism category, and specifically the aminobenzoate degradation pathway, generate catechol (1,2-dihydroxybenzene), which promotes Enterobacteriaceae growth and virulence [20]. This can explain the enrichment of this pathway (Fig S1A, S1B) and its positive correlation with Enterobacteriaceae, and may suggest a causative scenario: the change in the microbiota leading to Enterobacteriaceae enrichment. Taken together, the functional prediction of the microbiota leading to enrichment in Enterobacteriaceae, immune system modulation and intestinal epithelial damage, can explain the higher rate of blood-stream infections in the CRE-carrierssince colonization with Enterobacteriaciae has been associated with increased risk for bacteremia [21].

These compositional and functional changes predispose the host to invasive infection and death. We found higher rates of bacteremia (not caused only by CRE) in the CRE-carriers group compared to the non-carriers (Table 1). Interestingly, only one patient had bacteremia with *K. pneumoniae* KPC, which is consistent with a previous study showing that *K. pneumoniae* isolates from blood samples were less likely to harbor KPC [3]. This can be explained by the fitness costs of resistance, typically observed as a reduced bacterial growth rate [22].

It can be assumed that the dysbiotic microbiota and high rate of bacteremia in CRE-carriers is linked by low levels of SCFAs, shown to interact with innate mechanisms of defense against infection (regulation of immune cell function by SCFA), and low levels of “defensive bacteria” such as *Barnesiella* [6,16].

Once established, the gut microbiota composition is relatively stable throughout adult life, but can alter as a result of the action of several vectors. In our study, a trend towards a statistically significant difference between the experimental groups was found in the following factors: treatment with carbapenem, chemotherapy treatment, and gastrointestinal disease/disorder. However, we cannot point to the exact determinants influencing the microbiota composition in CRE-carriers. Whatever the predominant factors that modify the microbiota are, the end result is an “unhealthy microbiota” which lost “key species” required for shaping a “healthy microbiota.” Indeed, gut microbiota has been previously shown to effect susceptibility to infections caused by other pathogens such as *Vibrio cholerae* [23], and *C. difficile* [24].

One major limitation of our study is the strong impact of antibiotic treatment on the gut microbiota [25]. To weaken the effect, one of our control groups was comprised of hospitalized non-CRE-carriers. Both hospitalized groups were hospitalized (for at least seven days) at the same healthcare facility, and were treated with antibiotics profile similar to that of the hospitalized CRE-carriers. As mentioned, no significant confounding factors differed between the CRE-carriers and non-carriers, except for the CRE-carriage itself. There may be other unexamined factors that may cause the microbial differences we found. Moreover, all taxa-associated analyses were conducted by excluding the healthy controls (not hospitalized), since the lack of antibiotic treatment does not allow proper comparison between the groups.

In addition, on the basis of this study it is not possible to determine causality between dysbiosis and CRE-colonization, as dysbiosis can be both a cause and a result of CRE-colonization.

## Conclusions

Overall the results in our cohort, indicate that the interrelation between dysbiotic microbiota, its pool of bacterial genes (microbiome), and their expressing functions might weakens the protection and resistance against CRE-colonization and infection and other pathobionts. Therefore, our study supports the possibility of fecal transplantation as a therapeutic strategy for CRE-carriage, a strategy already efficiently used to treat *C. difficile* recurrent infection [16,26] and studied for eradication of other highly drug-resistant enteric bacteria carriage [7,27]. Reintroducing specific strains and/or correction of dysbiosis with probiotics or fecal transplantation may potentially lead to colonization resistance restoration [7,27].

## References

1. Snitkin ES, Zelazny AM, Thomas PJ, Stock F, NISC Comparative Sequencing Program Group NCS, Henderson DK, et al. Tracking a hospital outbreak of carbapenem-resistant Klebsiella pneumoniae with whole-genome sequencing. Sci Transl Med. 2012;4:116–48.

2. Gupta N, Limbago BM, Patel JB, Kallen AJ. Carbapenem-Resistant Enterobacteriaceae: Epidemiology and Prevention. Clin Infect Dis [Internet]. Oxford University Press; 2011 [cited 2018 Oct 28];53:60–7. Available from: https://academic.oup.com/cid/article-lookup/doi/10.1093/cid/cir202

3. Gasink LB, Edelstein PH, Lautenbach E, Synnestvedt M, Fishman NO. Risk factors and clinical impact of Klebsiella pneumoniae carbapenemase-producing K. pneumoniae. Infect Control Hosp Epidemiol. 2009;30:1180–5.

4. Campos AC, Albiero J, Ecker AB, Kuroda CM, Meirelles LEF, Polato A, et al. Outbreak of Klebsiella pneumoniae carbapenemase–producing K pneumoniae: A systematic review. Am J Infect Control [Internet]. Mosby; 2016 [cited 2018 Oct 28];44:1374–80. Available from: https://www.sciencedirect.com/science/article/pii/S0196655316002807?via%3Dihub

5. Bilavsky E, Schwaber MJ, Carmeli Y. How to stem the tide of carbapenemase-producing enterobacteriaceae?: proactive versus reactive strategies. Curr Opin Infect Dis. 2010;23:327–31.

6. Montassier E, Al-Ghalith GA, Ward T, Corvec S, Gastinne T, Potel G, et al. Pretreatment gut microbiome predicts chemotherapy-related bloodstream infection. Genome Med [Internet]. 2016;8. Available from: http://genomemedicine.biomedcentral.com/articles/10.1186/s13073-016-0321-0

7. Caballero S, Kim S, Carter RA, Leiner IM, Sušac B, Miller L, et al. Cooperating Commensals Restore Colonization Resistance to Vancomycin-Resistant Enterococcus faecium. Cell Host Microbe [Internet]. Cell Press; 2017 [cited 2017 Dec 4];21:592–602.e4. Available from: https://www.sciencedirect.com/science/article/pii/S1931312817301488

8. Ubeda C, Taur Y, Jenq RR, Equinda MJ, Son T, Samstein M, et al. Vancomycin-resistant Enterococcus domination of intestinal microbiota is enabled by antibiotic treatment in mice and precedes bloodstream invasion in humans. J Clin Invest. 2010;120:4332–41.

9. Taur Y, Xavier JB, Lipuma L, Ubeda C, Goldberg J, Gobourne A, et al. Intestinal Domination and the Risk of Bacteremia in Patients Undergoing Allogeneic Hematopoietic Stem Cell Transplantation. CID. 2012;55:905–14.

10. Livorsi DJ, Arif S, Garry P, Kundu MG, Satola SW, Davis TH, et al. Methicillin-Resistant Staphylococcus aureus (MRSA) Nasal Real-Time PCR: A Predictive Tool for Contamination of the Hospital Environment. Infect Control Hosp Epidemiol [Internet]. 2015 [cited 2019 Mar 6];36:34–9. Available from: http://cran.us.r-project.org

11. Donskey CJ, Chowdhry TK, Hecker MT, Hoyen CK, Hanrahan JA, Hujer AM, et al. Effect of Antibiotic Therapy on the Density of Vancomycin-Resistant Enterococci in the Stool of Colonized Patients. N Engl J Med [Internet]. Massachusetts Medical Society; 2000 [cited 2019 Mar 6];343:1925–32. Available from: http://www.nejm.org/doi/abs/10.1056/NEJM200012283432604

12. Ríos-Covián D, Ruas-Madiedo P, Margolles A, Gueimonde M, de Los Reyes-Gavilán CG, Salazar N. Intestinal Short Chain Fatty Acids and their Link with Diet and Human Health. Front Microbiol [Internet]. Frontiers Media SA; 2016;7:185. Available from: http://www.ncbi.nlm.nih.gov/pubmed/26925050

13. Shen P, Whelan FJ, Schenck LP, McGrath JJC, Vanderstocken G, Bowdish DME, et al. Streptococcus pneumoniae colonization is required to alter the nasal microbiota in cigarette smokeexposed mice. Infect Immun. 2017;85.

14. Integrative HMP Research Network Consortium T. The Integrative Human Microbiome Project: Dynamic Analysis of Microbiome-Host Omics Profiles during Periods of Human Health and Disease. Cell Host Microbe [Internet]. 2014;16:276–89. Available from: http://dx.doi.org/10.1016/j.chom.2014.08.014ThisisanopenaccessarticleundertheCCBYlicense

15. Sommer F, Anderson JM, Bharti R, Raes J, Rosenstiel P. The resilience of the intestinal microbiota influences health and disease. Nat Rev Microbiol [Internet]. Nature Publishing Group; 2017 [cited 2018 Oct 22];15:630–8. Available from: http://www.nature.com/doifinder/10.1038/nrmicro.2017.58

16. Maslowski KM, Mackay CR. Diet, gut microbiota and and immune responses. Nat Immunol. Nature Publishing Group; 2011;12:5–9.

17. Jiang X, Chen ZJ. The role of ubiquitylation in immune defence and pathogen evasion. Nat. Rev. Immunol. 2012. p. 35–48.

18. Ashida H, Kim M, Sasakawa C. Exploitation of the host ubiquitin system by human bacterial pathogens. Nat. Rev. Microbiol. 2014. p. 399–413.

19. Dinh DM, Volpe GE, Duffalo C, Bhalchandra S, Tai AK, Kane A V., et al. Intestinal Microbiota, microbial translocation, and systemic inflammation in chronic HIV infection. J Infect Dis. 2015. p. 19–27.

20. Rooks MG, Veiga P, Wardwell-Scott LH, Tickle T, Segata N, Michaud M, et al. Gut microbiome composition and function in experimental colitis during active disease and treatment-induced remission. ISME J [Internet]. 2014 [cited 2018 Jan 9];8:1403–17. Available from: https://www.nature.com/articles/ismej20143.pdf

21. Ruppé E, Andremont A. Causes, consequences, and perspectives in the variations of intestinal density of colonization of multidrug-resistant enterobacteria. Front Microbiol [Internet]. Frontiers; 2013 [cited 2019 Apr 7];4:129. Available from: http://journal.frontiersin.org/article/10.3389/fmicb.2013.00129/abstract

22. Andersson DI, Hughes D. Antibiotic resistance and its cost: is it possible to reverse resistance? 2010; Available from: https://www.nature.com/articles/nrmicro2319.pdf

23. Weil A, Midani F, Chowdhury F, Khan A, Begum Y, Charles R, et al. The Gut Microbiome and Susceptibility to Vibrio cholerae Infection. Open Forum Infect Dis [Internet]. Oxford University Press; 2015 [cited 2018 May 28];2. Available from: https://academic.oup.com/ofid/article-lookup/doi/10.1093/ofid/ofv131.72

24. Theriot CM, Koenigsknecht MJ, Carlson PE, Hatton GE, Nelson AM, Li B, et al. Antibiotic-induced shifts in the mouse gut microbiome and metabolome increase susceptibility to Clostridium difficile infection. Nat Commun [Internet]. Nature Publishing Group; 2014 [cited 2018 May 28];5:3114. Available from: http://www.nature.com/articles/ncomms4114

25. Jeffery IB, Lynch DB, O’Toole PW. Composition and temporal stability of the gut microbiota in older persons. ISME J. 2015;1–13.

26. Borody TJ, Khoruts A. Fecal microbiota transplantation and emerging applications. Nat Rev Gastroenterol Hepatol. 2011;9:88–96.

27. Manges AR, Steiner TS, Wright AJ. Fecal microbiota transplantation for the intestinal decolonization of extensively antimicrobial-resistant opportunistic pathogens: a review. Infect Dis (Auckl) [Internet]. 2016 [cited 2017 Dec 4];48:587–92. Available from: http://www.tandfonline.com/doi/full/10.1080/23744235.2016.1177199

28. Caporaso JG, Kuczynski J, Stombaugh J, Bittinger K, Bushman FD, Costello EK, et al. QIIME allows analysis of high-throughput community sequencing data. Nat Methods [Internet]. 2010;7:335–6. Available from: http://www.ncbi.nlm.nih.gov/pubmed/20383131

29. Segata N, Izard J, Waldron L, Gevers D, Miropolsky L, Garrett WS, et al. Metagenomic biomarker discovery and explanation. Genome Biol [Internet]. 2011;12:R60. Available from: http://www.ncbi.nlm.nih.gov/pubmed/21702898

30. Kanehisa M, Goto S. KEGG: kyoto encyclopedia of genes and genomes. Nucleic Acids Res [Internet]. 2000 [cited 2018 Aug 15];28:27–30. Available from: http://www.ncbi.nlm.nih.gov/pubmed/10592173

31. I Langille MG, Zaneveld J, Gregory Caporaso J, McDonald D, Knights D, Reyes JA, et al. Predictive functional profiling of microbial communities using 16S rRNA marker gene sequences. Nat Biotechnol [Internet]. 2013 [cited 2017 Nov 27];31. Available from: https://www.nature.com/articles/nbt.2676.pdf

